# ChEC-seq2: an improved Chromatin Endogenous Cleavage sequencing method and bioinformatic analysis pipeline for mapping *in vivo* protein-DNA interactions

**DOI:** 10.1101/2023.10.15.562421

**Authors:** Jake VanBelzen, Chengzhe Duan, Donna Garvey Brickner, Jason H. Brickner

## Abstract

Defining the *in vivo* DNA binding specificity of transcription factors (TFs) has relied nearly exclusively on chromatin immunoprecipitation (ChIP). While ChIP reveals TF binding patterns, its resolution is low. Higher resolution methods employing nucleases such as ChIP-exo, chromatin endogenous cleavage (ChEC-seq) and CUT&RUN resolve both TF occupancy and binding site protection. ChEC-seq, in which an endogenous TF is fused to micrococcal nuclease, requires neither fixation nor antibodies. However, the specificity of DNA cleavage during ChEC has been suggested to be lower than the specificity of the peaks identified by ChIP or ChIP-exo, perhaps reflecting non-specific binding of transcription factors to DNA. We have simplified the ChEC-seq protocol to minimize nuclease digestion while increasing the yield of cleaved DNA. ChEC-seq2 cleavage patterns were highly reproducible between replicates and with published ChEC-seq data. Combined with DoubleChEC, a new bioinformatic pipeline that removes non-specific cleavage sites, ChEC-seq2 identified high-confidence cleavage sites for three different yeast TFs that are strongly enriched for their known binding sites and adjacent to known target genes.

## Introduction

Chromatin immunoprecipitation is a powerful and widely used method that captures *in vivo* protein-DNA interactions through formaldehyde fixation and immunoprecipitation (Gilmour and Lis, 1984; Solomon et al., 1988). When combined with next generation sequencing (*i.e.* ChIP-seq), this method provides an unbiased approach to identifying sites genome-wide and, in the case of sequence-specific DNA binding proteins, to define their specificity (Johnson et al., 2007). However, the resolution of ChIP-seq is limited by the length of the isolated DNA fragments - typically hundreds of base pairs long and much longer than the size of transcription factor binding sites. An alternative approach, ChIP-exo, whereby formaldehyde crosslinked chromatin is digested with exonucleases to reveal protected regions, provides much higher resolution (Rhee and Pugh, 2011). Likewise, the CUT&RUN method, whereby Protein A is fused to micrococcal nuclease (MNase) and chromatin from purified nuclei is treated with a primary antibody against the factor of interest and Protein A-MNase to digest nearby DNA, followed by immunoprecipitation, greatly improves the resolution of ChIP-seq (Skene and Henikoff, 2017). However, these latter methods are technically challenging, produce small amounts of material and are dependent on formaldehyde fixation and antibody quality.

An alternative method to map protein-DNA interactions *in vivo* is Chromatin Endogenous Cleavage (ChEC; Zentner et al., 2015; Schmid et al., 2004), which utilizes micrococcal nuclease (MNase) fused to the protein of interest. MNase activity is maximal at 10 mM Ca^2+^, far greater than the ∼150nM concentration found in the nucleus of budding yeast (Ohya et al., 1991), so cells can tolerate the expression of MNase fusion proteins. However, upon permeabilization of cells in the presence of millimolar calcium, MNase rapidly cleaves DNA nearby (Schmid et al., 2004). This method was adapted for next generation sequencing (ChEC-seq; Zentner et al., 2015) by isolating small DNA fragments through negative selection with AMPure Beads, repairing DNA ends, and ligation of Illumina TruSeq adapters for sequencing (Figure 1a). Small DNA fragments arise from DNA that was cleaved at two adjacent sites, which are enriched for the binding site of the protein of interest (Zentner et al., 2015). However, these small DNA fragments are low in abundance and therefore difficult to reproducibly purify, and their abundance is strongly influenced by the number of binding sites and the concentration of the protein of interest. Proteins that sparsely interact with the genome yield relatively few small DNA fragments, making ChEC-seq less feasible. Extending the cleavage time seems to increase non-specific cleavage rather than increasing the yield of specifically cleaved fragments (Zentner et al., 2015).

**Figure 1.**
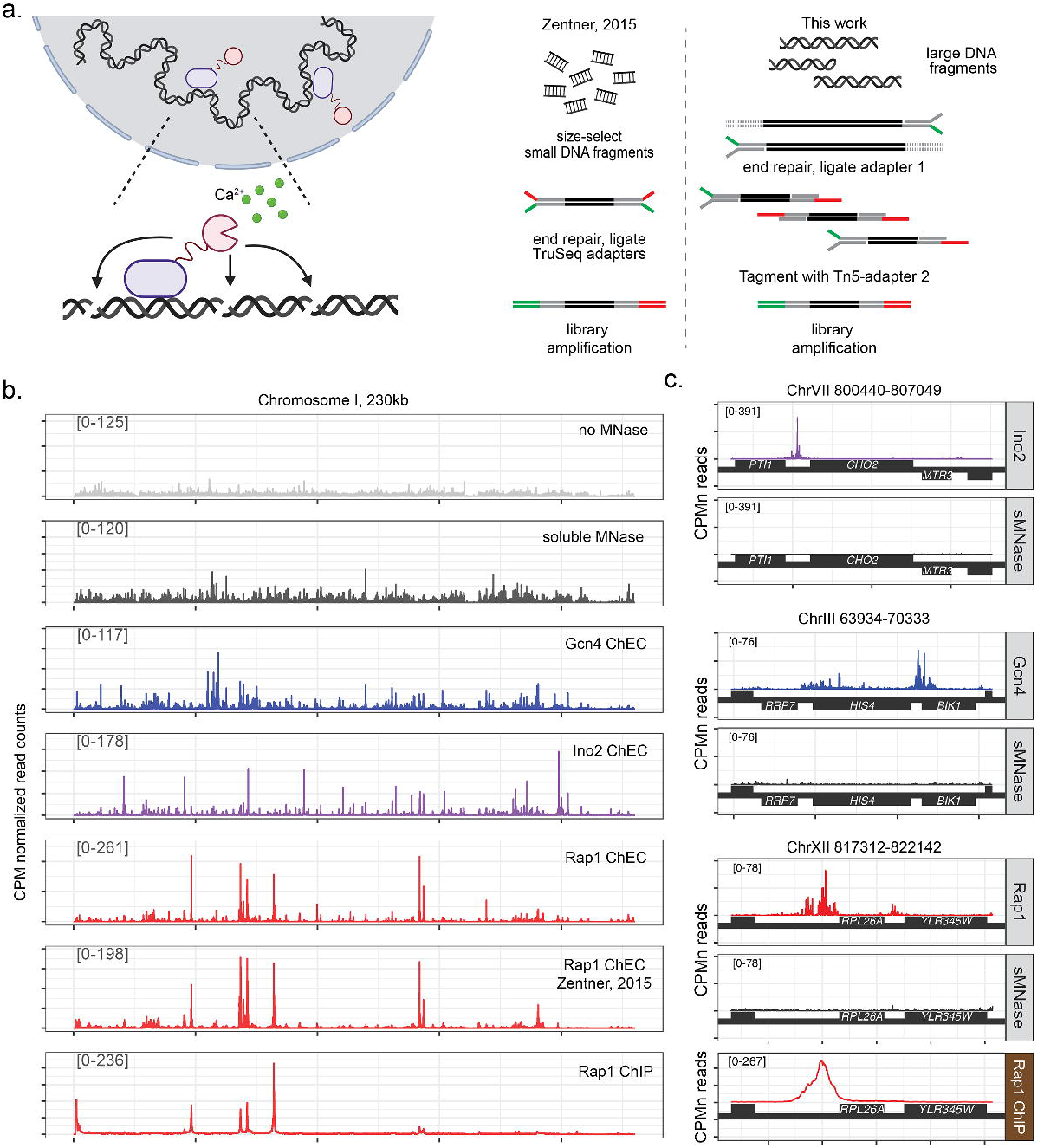
An improved method for chromatin endogenous cleavage (ChEC-seq2). **(a)** Schematic of modifications of ChEC-seq method. Left: a DNA-bound protein fused with micrococcal nuclease (MNase; pink). Upon permeabilization of the cells and addition of calcium, MNase cleaves adjacent DNA. Right: whereas the original method requires size-selection of small quantities of small fragments that have been cleaved twice, the current work uses mild digestion to generate large, cleaved fragments (Supplementary Figure 1). These large fragments are repaired and ligated to Read adapter 1 (green). Treatment with Tn5 transposase loaded with Read adapter 2 (red) results in small fragments that are amplified for single-end sequencing from adapter 1. **(b)** CPM-normalized read coverage from ChEC-seq experiments with Gcn4, Ino2, Rap1, soluble MNase (sMNase), and no MNase over *S. cerevisiae* Chromosome 1. The average of 3-biological replicates is shown. For ChEC-seq data, mapped reads were trimmed to the first base pair adjacent to the cleaved DNA. Published Rap1 ChEC-seq and Rap1 ChIP-seq datasets are included for comparison in the bottom two panels. **(c)** Representative targets of each transcription factor displayed as in **(b)** with soluble MNase (grey) for comparison, along with Rap1 ChIP-seq data (bottom panel).

To address this issue, we have developed ChEC-seq2, which accepts all of the digested genomic DNA as input, marks free ends through the ligation of a custom adapter, followed by Tn5 transposase-mediated library construction (Figure 1a). Library amplification with Nextera index primers amplifies DNA fragments flanked by heterologous adapters, and the final product is a DNA library compatible with Illumina sequencing. This method generates a much higher yield of DNA, minimizing the amplification needed to generate the library. This is particularly true for transcription factors that bind fewer sites. The cleavage patterns are highly reproducible and agree well with data from the original ChEC-seq method.

A significant concern with ChEC-seq is the specificity of the cleavage pattern. ChEC-seq with transcription factors (TFs) identifies cleavage both at specific sites with recognizable motifs as well as many more non-specific sites (Zentner et al., 2015), although cleavage of specific sites was observed preferentially at shorter cleavage times. This was interpreted to represent cleavage by both specifically bound TFs and non-specifically scanning TFs, although that conclusion has been challenged (Rossi et al., 2017). Furthermore, non-specific cleavage of unprotected DNA is a source of background that must be accounted for to interpret ChEC data (Mittal et al., 2021). We have developed a bioinformatic filtering approach called DoubleChEC that identifies high-confidence targets based on 1) the enrichment of cleavage compared with a negative control that evaluates chromatin accessibility, 2) the structure of the cleavage pattern and 3) the protection of the binding site. This filtering produces very robust identification of TF binding motifs and target genes and is not affected by the abundance or type of transcription factor. ChEC-seq2 with three different TFs identified known binding sites and target genes and agrees well with published ChIP-seq and ChIP-exo datasets. This method and bioinformatic pipeline provide a simple and robust method for mapping protein-DNA interactions *in vivo*.

## Results

A detailed ChEC-seq2 protocol is provided as a supplementary file. Partial MNase digestion was optimized for each fusion protein to produce large fragments (> 5kb; mean size ∼ 15kb; supplementary Figures 1a-c). DNA was purified from cells, cleaved ends were repaired and ligated to an P5-compatible Y-adapter (containing the Read 1 adapter sequence; Figure 1a). Repaired DNA fragments were then fragmented with recombinant Tn5 transposase (Hennig et al., 2017) loaded with Read 2 adapter (*i.e.* “Tagmented”, Figure 1a). Similar to the Tagmentation in Nextera Library Preparation, this yielded a mixture of small DNA fragments flanked by adapter sequences (< 1kb; Supplementary Figure 1d). DNA fragments bearing both Read 1 and Read 2 adapter regions were enriched by 15 cycles of PCR amplification using Nextera XT primers. The indexed libraries were pooled and sequenced using Illumina HiSeq 4000 single-end 50bp reads. The reads are from the Read 1 direction, which represents the bases adjacent to the cleavage site.

### ChEC-seq2 is highly reproducible and recapitulates the patterns produced by ChEC-seq

To evaluate ChEC-seq2, we fused MNase to three well-characterized budding yeast transcription factors, Rap1, Gcn4, and Ino2 at their endogenous loci. Rap1, Gcn4 and Ino2 represent a good test of the robustness of ChEC-seq2 because they bind to distinct, well-defined sequences (https://jaspar.genereg.net/; Figure S1; Castro-Mondragon et al., 2021) and have well-understood biological functions. Furthermore, they differ in abundance (Rap1 ∼ 4400 molecules per cell; Gcn4 <100 molecules per cell; Ino2 ∼ 784 molecules per cell; (Ghaemmaghami et al., 2003) and regulate different numbers of target genes (Rap1, 906 targets; Gcn4, 1257 targets, Ino2, 86 targets; Saccharomyces Genome Database). ChEC-seq2 was also performed with a strain expressing soluble, nuclear MNase (sMNase) expressed from the *CYC1* promoter to control for chromatin accessibility and the sequence bias of MNase digestion (Horz et al., 1981). Each TF cleaved at a different rate, likely reflecting its abundance and number of DNA-binding sites. Time points that produce clear, partial digestion were selected for both the TFs and sMNase from cells grown under identical conditions (Figure S2).

The first base of each sequencing read corresponds to the genomic location adjacent to a MNase cleavage event. Therefore, following read-mapping, genome-coverage data was calculated from only the first base of each read, which facilitates resolution of the fine structure of the cleavage peaks (Figure S3). Genome coverage data was then normalized to the library size (counts per million reads, CPM) and averaged from three biological replicates. The pattern of cleavage across chromosome I revealed patterns for each transcription factor (Figure 1b) that were distinct from each other and from libraries made from a strain lacking MNase (representing mechanical shearing) or expressing sMNase (Figure 1b). Comparison to previously published Rap1 ChEC-seq data revealed that ChEC-seq2 produces a highly similar cleavage pattern (Figure 1b). A high-quality Rap1 ChIP-seq dataset (Bondra and Rine, 2023) showed a similar, but not identical, pattern of peaks to the ChEC-seq or ChEC-seq2 datasets (Figure 1b). Finally, the peaks of cleavage from ChEC are much narrower than those produced by ChIP-seq (Figure 1c).

### DoubleChEC: a filtering approach to identify high-confidence transcription factor binding sites

ChEC-seq and ChEC-seq2 reveal cleavage peaks of different intensity and only a fraction of these sites are *bona fide* binding sites that resemble the consensus motif (Zentner et al., 2015). The biological significance of such “off-target” cleavage events is unclear. They may represent transcription factor scanning or chromatin accessibility to randomly diffusing DNA binding proteins. Regardless, identifying sites with high occupancy is critical to define the sequence specificity and biological role of DNA binding proteins. Previous work distinguished between high occupancy and low occupancy sites based on relative peak height and how rapidly the peaks appeared in time course experiments (Zentner et al., 2015). We reasoned that true binding sites should be associated with peaks that are significantly larger than those produced by soluble MNase, which controls for DNA accessibility and MNase bias (Horz et al., 1981), and they should be found in pairs, flanking protected binding sites. To test this hypothesis, we measured the mean Rap1-MN or sMNase cleavage patterns over putative Rap1 binding sites (2366 instances of 5’-GNNNGGGTG-3’; Figure 2a). Indeed, Rap1-MN cleavage frequency produced strong peaks immediately adjacent to the binding site with a protected region of 8-10 bp over the query sequence (Figure 2a). The cleavage frequency of sMNase was generally low and also reflected protection of the query sequence (Figure 2a). Thus, *bona fide* TF binding sites should produce pairs of peaks flanking a protected sequence.

**Figure 2.**
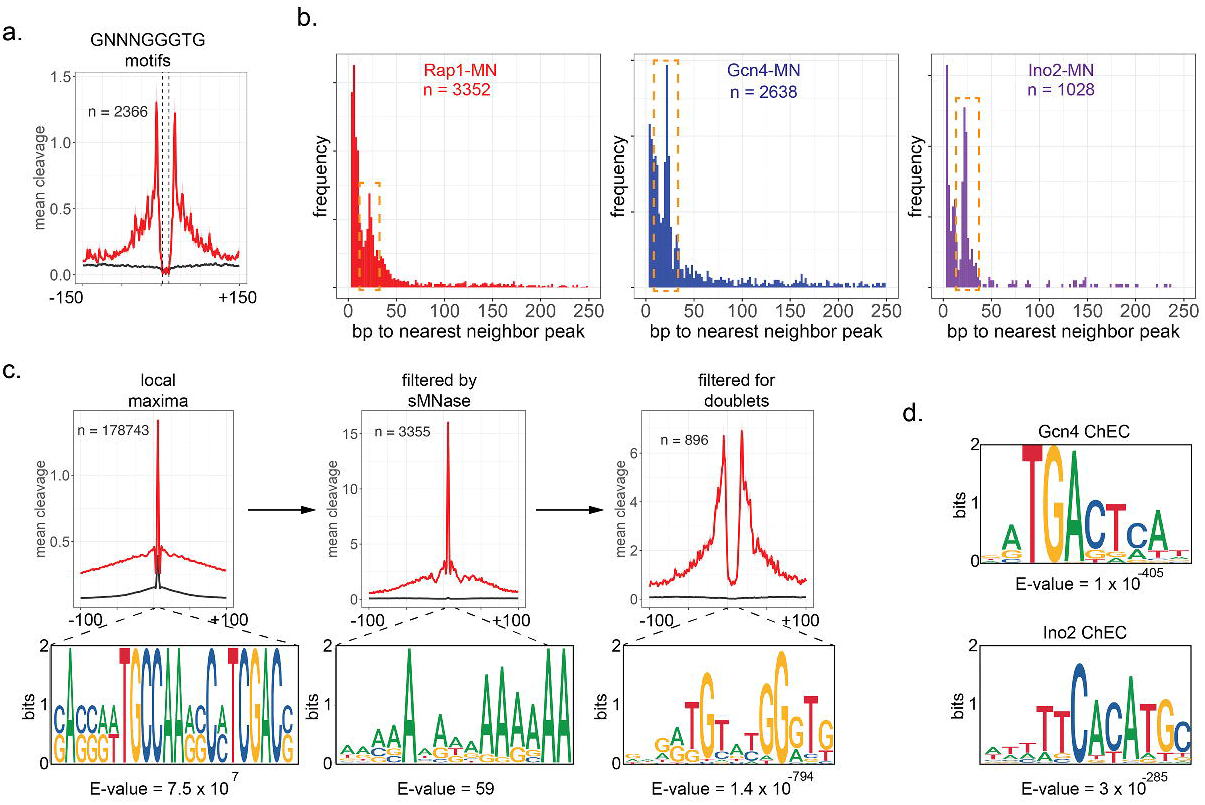
Improved analysis pipeline to identify high-confidence binding sites from ChEC-seq2 data. **(a)** Mean Rap1 ChEC-seq2 cleavage (red) and soluble MNase cleavage (black) over all 5’-GNNNGGGTG-3’ sites in the yeast genome. The average of 3 biological replicates is displayed. The dashed lines flank an 8bp protected region over the sequence. **(b)** Distribution of distances between adjacent peaks that were significantly enriched over sMNase. A secondary peak between 15bp and 50bp is highlighted with the dashed box. **(c)** Mean cleavage by Rap1 (red) and soluble MNase (black) over local maxima (left), peaks enriched over soluble MNase (middle) or for doublets (right). Each set of peaks (21bp windows; overlapping peaks were merged) or peak pairs (merged) was analyzed for motif enrichment using MEME and the most enriched motif from each set is shown below. **(d)** High confidence sites for Gcn4 and Ino2 from ChEC-seq2 data were identified as in **(b).** Doublet peaks were analyzed by MEME and the top motifs for each Gcn4 (top) and Ino2 (bottom) are shown.

To identify high-confidence binding sites, we developed a workflow in R (https://www.r-project.org/) called DoubleChEC that: 1) smooths the ChEC cleavage frequency using a sliding window of 3bp, step size of 2bp, 2) identifies all local maxima, 3) compares the cleavage of the protein of interest to sMNase over every peak using DESeq2 (parameters: >1.7 log_2_-fold change, adjusted p-value < 1e-4; Love et al., 2014) and 4) identifies pairs of peaks. The optimal distance between pairs of peaks was determined by plotting the frequency of distances between adjacent TF peaks for Rap1, Gcn4 and Ino2. The distances between neighboring local maxima are dominated by peaks that are adjacent to each other (*i.e.* the peak is in the next neighboring sliding window; Figure S4a). However, for peaks that are significantly enriched over sMNase, a clear secondary maximum between 15bp and 50 bp is evident (Figure 2b). Therefore, we selected pairs of peaks with local maxima between 15 and 50bp apart (DoubleChEC is available at https://github.com/jasonbrickner/DoubleChEC.git).

To confirm that sMNase cleavage reflects non-specific cleavage, we also mapped the cleavage of chromatin by two chromatin-associated proteins, H2A.Z and Prp20. H2A.Z is enriched in nucleosomes near promoters (Raisner et al., 2005; Guillemette et al., 2005) and Prp20 is a general nucleosome binding protein (Frasch, 1991). The pattern of cleavage by H2A.Z and Prp20 was qualitatively very similar to that produced by sMNase (Supplementary Figure S5). Using either Prp20 or H2A.Z as a control instead of sMNase also identified strongly enriched motifs (Supplementary Figure S5). Therefore, sMNase or chromatin-associated proteins adequately control for non-specific cleavage in ChEC-seq2.

For Rap1-MN, 178,843 local maxima were identified from ChEC cleavage frequencies. Filtering for those that were significantly enriched over sMNase reduced this number to 3,352 peaks and increased average peak height by approximately 10-fold (Figure S4b & Figure 2c). Finally, selecting adjacent peaks flanking a protected region produced 896 doublet peaks (Figure 2c), each of which was merged into a single high-confidence site. MEME analysis (Bailey, 2003; Yu et al., 2012) revealed that neither the initial local maxima nor the sMNase filtered peaks were enriched for the Rap1 binding site (Figure 2c, bottom panels). However, doublet peaks gave a strong enrichment for the consensus Rap1 binding site (Figure 2c). Furthermore, this approach identified 627 doublet peaks for Gcn4 (during histidine starvation) and 194 doublet peaks for Ino2 (during inositol starvation) that were strongly enriched for their respective consensus binding sites (Figure 2d).

To assess the ability of DoubleChEC to identify target genes, genes with high-confidence sites identified by ChEC-seq2 within 700 bp upstream were identified for Rap1-MN (691 genes), Gcn4-MN (487 genes) and Ino2-MN (203 genes). Gene ontology analysis (Ashburner et al., 2000) of these genes revealed that the top 10 most significantly enriched terms for each set matched the known biological functions of Rap1 (regulator of ribosome biosynthesis), Gcn4 (regulator of amino acid biosynthesis) and Ino2 (regulator of phospholipid biosynthesis). These genes were also compared with TF targets reported for 133 transcription factors by the Saccharomyces Genome Database (SGD; https://yeastmine.yeastgenome.org/yeastmine/begin.do; downloaded 09-25-2023; Supplementary Table S1; Balakrishnan et al., 2012). Enrichment was assessed by Fisher’s Exact test (Bonferroni corrected p-value). For Rap1-MN and Gcn4-MN, the overlap was most significant with their respective target sets from SGD (Figure 3b). Furthermore, Rap1-MN ChEC-seq2 target genes were strongly enriched for Fhl1 and Ifh1, transcription factors that co-regulate Rap1 targets (Shore et al., 2021). In contrast, Ino2 targets were only modestly enriched among the genes near Ino2-MN ChEC-seq2 sites (adjusted p-value = 0.03). However, the genes near Ino2-MN sites were enriched for targets of Ino4 (with which Ino2 heterodimerizes; adjusted p-value = 3×10^-3^) and Opi1 (to which Ino2 binds directly; adjusted p-value = 7×10^-4^; (Wagner et al., 2001; Heyken et al., 2005). This suggests that the list of 86 targets on SGD is incomplete. Indeed, comparison of genes near Ino2-MN sites with genes near ChIP-exo peaks (Bergenholm et al., 2018), revealed highly significant overlap (adjusted p-value = 4×10^-9^; labeled INO2* in Figure 3b). Thus, ChEC-seq2 is an efficient and robust method for identifying TF binding sites, sequence motifs and target genes.

**Figure 3.**
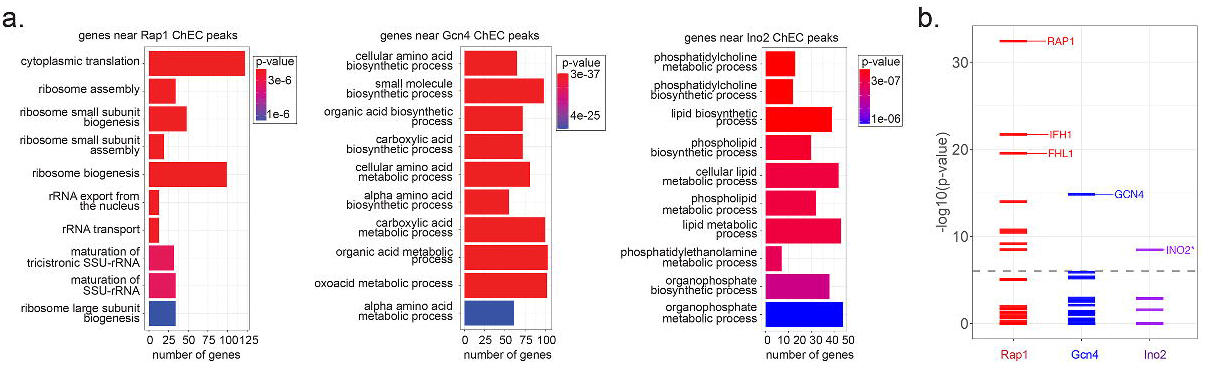
Target gene enrichment. **(a)** Genes with high-confidence sites from Rap1-MN (left; 691 genes), Gcn4-MN (middle; 487 genes), and Ino2-MN (right; 527) ChEC-seq2 within 700bp upstream of their start codon were analyzed for gene ontology (GO) term enrichment. The top 10 GO terms and their adjusted p-values are shown. **(b)** Genes adjacent to high-confidence sites from Rap1-MN, Gcn4-MN or Ino2-MN ChECseq2 were compared with reported target genes from 133 yeast transcription factors (www.yeastgenome.org). The significance of the overlap was assessed using a Fisher Exact test and the -log_10_(Bonferroni-adjusted p-value) for each comparison was plotted. The dashed line represents an adjusted p-value of 1 x 10^-6^. The identities of the top three TFs with an adjusted p-value < 1 x 10^-6^ is shown.

Unlike Rap1, Gcn4 and Ino2 function conditionally. Gcn4 protein levels are upregulated by starvation for amino acids (Hinnebusch, 1985). Ino2 is regulated by a repressor protein (Opi1), which dissociates upon starvation for inositol (Wagner et al., 2001; Heyken et al., 2005). However, Ino2 expression is also induced by inositol starvation, and the occupancy of Ino2 increases under activating conditions (Brickner and Walter, 2004). The cleavage over high-confidence Gcn4 and Ino2 sites increased under inducing conditions, particularly for Gcn4 (Supplementary Figure S6). This suggests that TF occupancy impacts the amount of cleavage detected by ChEC-seq2.

### Comparison of ChEC-seq2 with other methods

The high-confidence peaks identified for Rap1-MN by ChEC-seq2 were first compared with the Rap1-MN peaks identified using the original ChEC-seq protocol (Zentner et al., 2015). The 7260 peaks identified from those data included nearly all of the 896 peaks identified by ChEC-seq2 (Figure 4a). However, the Rap1 motif (Figure 3a) was not enriched in this larger set of peaks, presumably because they include many non-specific peaks. When the data from 30 s of cleavage with Rap1-MN from Zentner et al., 2015 were filtered using the sMNase data presented here, followed by selection of doublet peaks, the non-overlapping peaks were largely removed (Figure 4b). The filtered set of 1667 high-confidence sites included 713 peaks from ChEC-seq2 reported here (*p* = 3.5 x 10^-132^, Fisher’s Exact test) and were strongly enriched for the Rap1 motif (Figure 4b). Thus, ChEC-seq and ChEC-seq2 produce highly concordant data and applying our analysis pipeline to published ChEC-seq data identifies high-confidence sites enriched for the Rap1 sequence motif.

**Figure 4.**
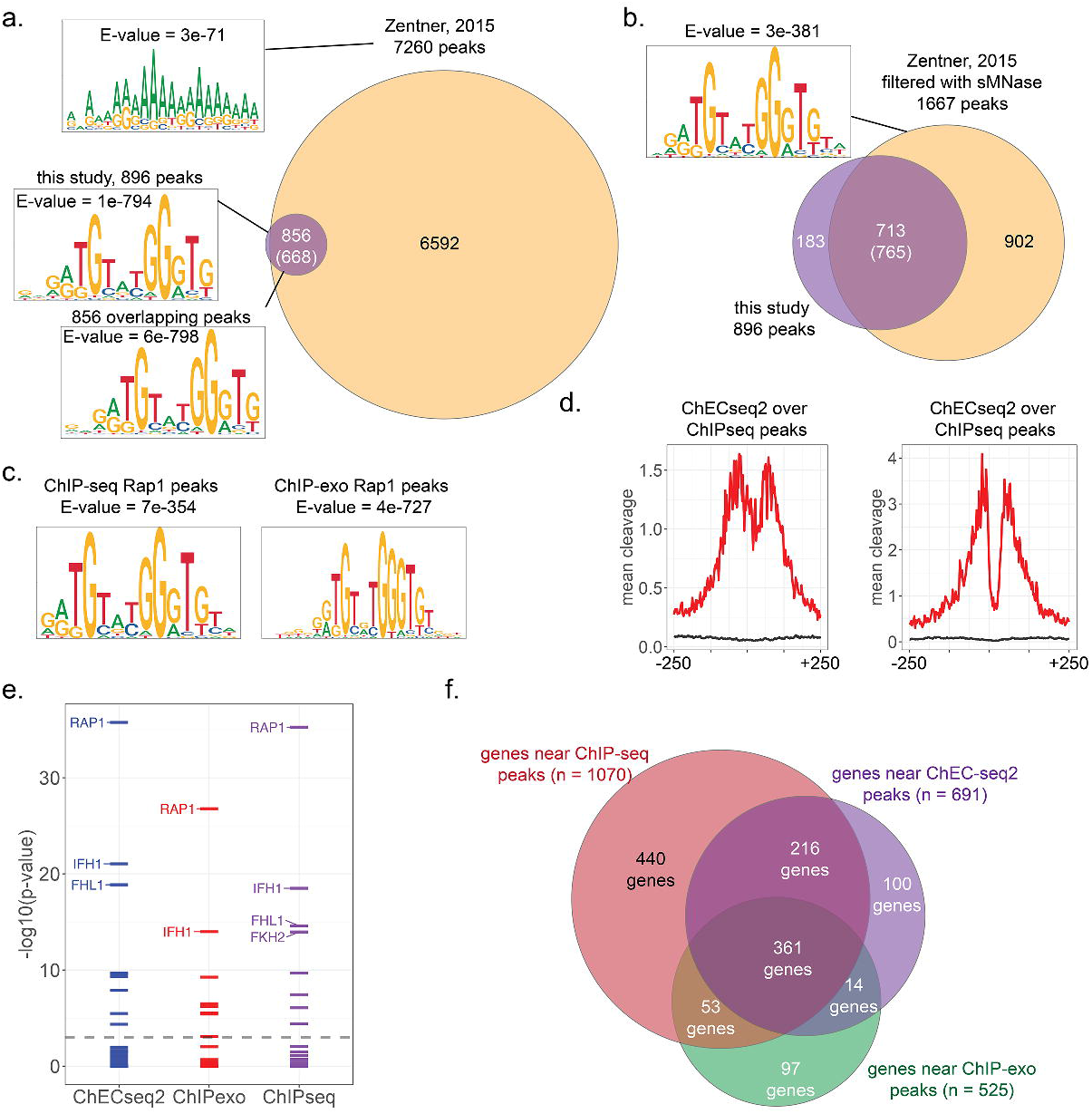
Comparison of ChEC-seq2 to other methods. **(a)** Overlap between 7260 Rap1 sites identified in Zentner et al., 2015 and 896 sites identified in the current work. Results of MEME analysis against all 7260, all 896 or the overlapping sites are shown. **(b)** High confidence Rap1 sites identified from 30s cleavage from Zentner et al., 2015 data, filtered using our peak finder (sMNase control from the current study). The top motif from the 1667 sites is shown. **(c)** Top motifs identified from Rap1 ChIP-seq and ChIP-exo sites. **(d)** Mean Rap1-MN cleavage over peaks identified by ChIP-seq and ChIP-exo. **(e)** Genes near ChEC-seq2, ChIP-seq and ChIP-exo peaks were compared for overlap with the targets of 133 TFs from SGD. The -log_10_(adjusted p-value) was plotted and the top three TFs with an adjusted p-value < 1e-6 were labeled. **(f)** Overlap between genes adjacent to ChEC-seq2 sites, ChIP-seq peaks and ChIP-exo peaks.

Finally, we compared the 896 high confidence Rap1 peak doublets identified by ChEC-seq2 with 1253 peaks identified by ChIP-seq (Bondra and Rine, 2023) or 576 peaks identified by ChIP-exo (Rhee and Pugh, 2011). The Rap1 motif is strongly enriched in all three sets of Rap1 sites (Figure 4c), although the degree of enrichment (*i.e.* E-value) was best for the ChEC-seq2 peaks (Figure 4a). Mean cleavage by Rap1-MN from our ChEC-seq2 data peaked over both sites identified by ChIP-seq and sites identified by ChIP-exo (Figure 4d). Genes near each of these sites (n = 691 for ChEC-seq2; n = 1070 for ChIP-seq and n = 525 for ChIP-exo) were compared for enrichment of the targets of 133 yeast TFs. All three sets showed strong enrichment for Rap1 as well as Fhl1/Ifh1 (Figure 4e). The target genes were strongly overlapping (Figure 4f). Of the 691 Rap1 targets identified by ChEC-seq2, 83% were also identified by ChIP-seq (*p* = 3 x 10^-61^, Fisher’s Exact test, Bonferroni corrected) and 54% were identified by ChIP-exo (*p* = 7 x 10^-80^).

## Discussion

Here we present a modified ChEC-seq protocol and analysis pipeline that provides excellent mapping of *in vivo* transcription factor binding sites. Similar to ChIP-seq and ChIP-exo, ChEC-seq2 identified consensus sequence motifs, target genes and biological roles for three different transcription factors in budding yeast that have distinct protein levels, regulation and DNA binding domains.

As an alternative to ChIP-seq or ChIP-exo, ChEC-seq2 has several advantages. ChEC-seq2 avoids fixation, which can affect chromatin solubility and may not be uniform for all binding sites. Also, ChEC-seq2 does not require antibody-based affinity purification, greatly increasing the yield of starting material and avoiding problems associated with variable accessibility of the epitope. Like ChIP-exo, ChEC-seq2 produces high resolution footprints of DNA binding, but unlike ChIP-exo, cleavage occurs in permeabilized cells and the downstream processing is simpler.

ChEC-seq2 has a disadvantage over ChIP-seq and ChIP-exo: non-specific cleavage of DNA at “off-target” sites throughout the genome. Such sites tend to be quite reproducible, enriched in nucleosome-depleted regions upstream and downstream of genes. Including these cleavage sites identifies a large number of non-biological sites (*i.e.* sites without an obvious DNA motif and that are not adjacent to known target genes). Selecting larger cleavage peaks or favoring sites that are produced more rapidly only partially addresses this issue. However, the cleavage pattern by soluble nuclear MNase or other chromatin-associated proteins controls for this artifact, allowing these sites to be removed. And together with selecting pairs of peaks flanking protected DNA, ChEC-seq2 identifies high-confidence sites enriched for consensus motifs.

While the sites and target genes identified by ChEC and ChIP are quite similar, some differences are apparent. The source of these differences may be biological (*i.e.* differences between strains or growth conditions) or technical (*i.e.* the ability to recover a particular site by ChIP or ChEC). It seems unlikely that variability of ChEC accounts for these differences; the agreement between ChEC replicates is very high. But it is possible that ChEC fails to recover some sites because of local DNA accessibility. Likewise, it is possible that differences in fixation efficiency or epitope availability could reduce recovery of particular sites by ChIP. The two techniques together seem to capture the most complete set of target sites for each transcription factor.

TFs represent an excellent test case for techniques such as ChIP and ChEC because they bind directly to well-defined DNA sequences in enhancer regions. However, many proteins occupy the genome less specifically. For example, co-activators or co-repressors frequently associate indirectly with enhancers through diverse TFs. This creates a challenge for cross-linking based methods because the antigen is unlikely to be directly cross-linked to DNA. Can ChEC-seq2 be used to map occupancy for such factors? It seems likely that such factors will enhance the cleavage of nearby DNA and that the significance of this cleavage can be assessed by comparison to sMNase. Although pairs of peaks are expected, the precise nature of the cleavage pattern, in terms of the distance between peaks and the degree of foot-printing, may differ from TFs. Therefore, it will be important to validate the analytical pipeline using genetics to ensure that binding sites are biologically meaningful.

## Methods

### Yeast strains

All yeast strains were derived from BY4741 and are listed in Supplementary Table 2. A C-terminal MNase fusion was introduced to the protein of interest through transformation and homologous recombination of PCR-amplified DNA. Primers were designed with 50-bp of homology to the 3’ end of the coding sequence of interest. The 3xFLAG-MNase with a KanR marker was amplified from pGZ108 (Zentner et al., 2015) and transformed into BY4741 as previously described. Successful transformation was confirmed by immunoblotting and PCR, followed by sequencing.

Lyophilized DNA oligonucleotides were resuspended in molecular-grade water to a concentration of 100 µM. For ligation, the following pair of oligonucleotides were annealed to produce the Y-adapter: Tn5ME-A (5’-TCGTCGGCAGCGTCAGATGTGTATAAGAGACAG-3’) and Y-Adapt-i5 R (5’-CTGTCTCTTATACACATCTTCATAGTAATCATC-3’). For Tn5 Tagmentation, the following i7 oligonucleotides were annealed: Tn5ME-B (5’ - GTCTCGTGGGCTCGGAGATGTGTATAAGAGACAG-3’) and Tn5MErev, (5’-PO_4_-CTGTCTCTTATACACATCT-3’). Pairs of oligonucleotides were annealed as follows: 45 µl of each oligo (100 µM) was combined with 10 µl of 1 M Potassium Acetate, 300 mM HEPES, pH 7.5 in a 0.2 ml PCR tube. In a thermocycler, the mixture was heated to 95<C for 4 minutes, cooled 1°C/minute until 50°C, incubated at 50°C for 5 minutes, and then cooled 1°C/minute until 4°C. Hybridized oligos were stored in 15 µl aliquots at -20<C.

### Tn5 purification and adapter loading

Tn5 E54K L372P was purified as previously described (Hennig et al., 2017). We found that Tn5 was sufficiently pure following purification on Ni^2+^-chromatography and we therefore omitted the final gel filtration step. Purified Tn5 was aliquoted and stored at -80°C. Optimal Tn5 activity was determined by cleaving genomic DNA and assessing fragmentation using the Femto Pulse (Figure S2d), and resulting DNA libraries were confirmed to be of appropriate length for Illumina Sequencing by TapeStation (Figure S2e).

Tn5 was thawed on ice and 100 µl Tn5 was added to 10 µl i7 (45 µM) in a 1.7 ml tube and mixed by gently pipetting. The mixture was incubated at 23°C, mixing at 350 rpm for 45 minutes. Adapter-loaded Tn5 was stored at -20°C and used within 24 hours.

### Chromatin endogenous cleavage and library preparation

Chromatin cleavage, DNA purification, end-repair, ligation, Tagmentation and library amplification was performed as described in the detailed protocol in the Supplementary Materials. Libraries were sequenced on an Illumina Hiseq and the Northwestern NUSeq core facility using the 50-bp, single-end option.

### Bioinformatic analysis

#### Quality Control, Trimming, and Mapping

Read quality and sequencer performance was evaluated with FASTQC. Reads were adapter and quality trimmed with Trimmomatic (Bolger et al., 2014) using single-end settings. Bases at either end of a read were trimmed if base-call quality was less than 30, and only reads of length ≥25 bp were retained. Trimmed reads were mapped to the *Saccharomyces cerevisiae* genome (Engel et al., 2013), version R64-4-1 with Bowtie2 (Langmead and Salzberg, 2012) and mapped reads with a MAPQ <10 were removed with Samtools (Li et al., 2009).

#### DoubleChEC identification of high-confidence TF binding sites

For peak calling analysis, BAM files for three or more biological replicates of the TF-MNase and soluble MNase were read and trimmed to the first base pair. Unnormalized counts and normalized counts per million (CPMn) were tallied for each base pair in the yeast genome and the average CPMn values among replicates were calculated for each position. Next, mean CPMn values were smoothed using a sliding window of 3 and a step size of 2. Windows with CPMn values less than three times the genome average were filtered out. After this filtering, local maxima (windows with values greater than their immediate neighbors) were identified. Unnormalized reads were smoothed, retaining positions that were identified as local maxima, and inputted them in DESeq2 (version 1.36.0) to identify windows with values significantly higher than those in the soluble MNase control. Only TF-MNase peaks with a greater log2-fold change of 1.7 and an adjusted p-value less than 0.0001 over soluble MNase were retained. Finally, the peaks were filtered again to identify doublet peaks that are between 15bp and 50bp apart, which were merged to single peaks.

#### GO term plot

A list of genes whose 700bp upstream regions overlap with peaks identified by the peak finder was input to enrichGO (Wu et al., 2021) to generate GO term plots based on biological functions. The 10 most significant GO terms with adjusted p-values less than 0.05 were plotted.

#### MEME analyses

The MEME Suite (version 5.5.1) was installed onto the local computer and two custom wrapper functions were written in R for the local bed2fasta and meme programs. These functions were then used to convert bed files, generated from peak calling, into FASTA files. These FASTA files were subsequently to generate motif logos. Both bed2fasta and meme programs were run using their default parameter values.

## Data availability

All sequencing data has been deposited at the NIH Short Read Archive (https://www.ncbi.nlm.nih.gov/sra/, BioProject #PRJNA1027910). The DoubleChEC peak finder R scripts are available at https://github.com/jasonbrickner/DoubleChEC.git).

## Supporting information

Strain table S2

Detailed ChEC-seq2 protocol

All TF targets

## Acknowledgements and author contributions

The authors thank Professor Erik Andersen, Professor Shelby Blythe and members of the Brickner laboratory for helpful comments on the manuscript. JV was supported by a National Science Foundation Graduate Fellowship. JV and DGB performed the experiments; JV, JHB and CD analyzed the data; JV and JHB wrote the manuscript. This work was supported by NIGMS R35GM136419 (JHB).

## Figure Legends

**Figure S1.**
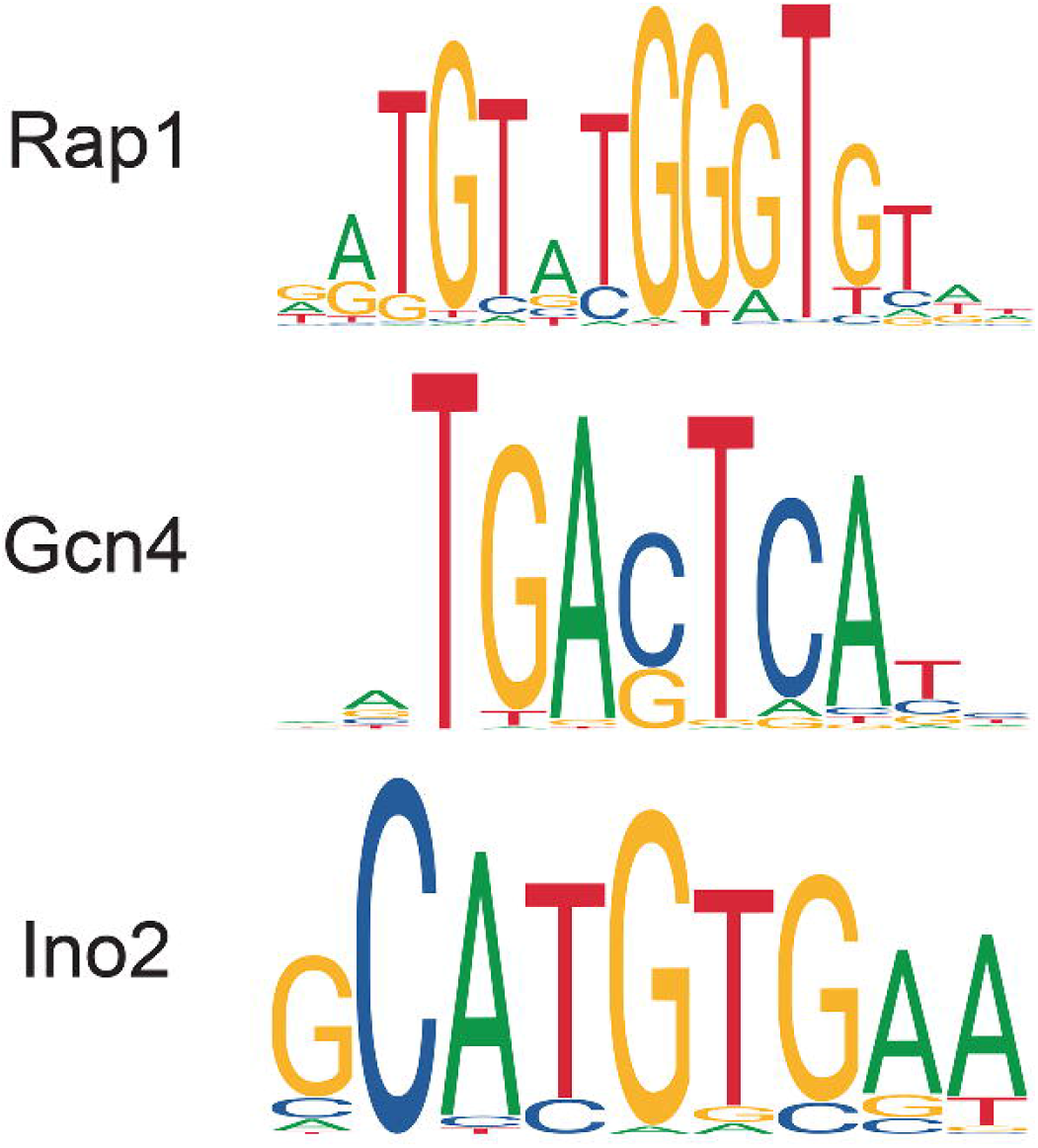
Consensus motifs for Rap1, Gcn4 and Ino2. Motifs for Rap1, Gcn4 and Ino2 from *Saccharomyces cerevisiae* from https://jaspar.genereg.net/.

**Figure S2.**
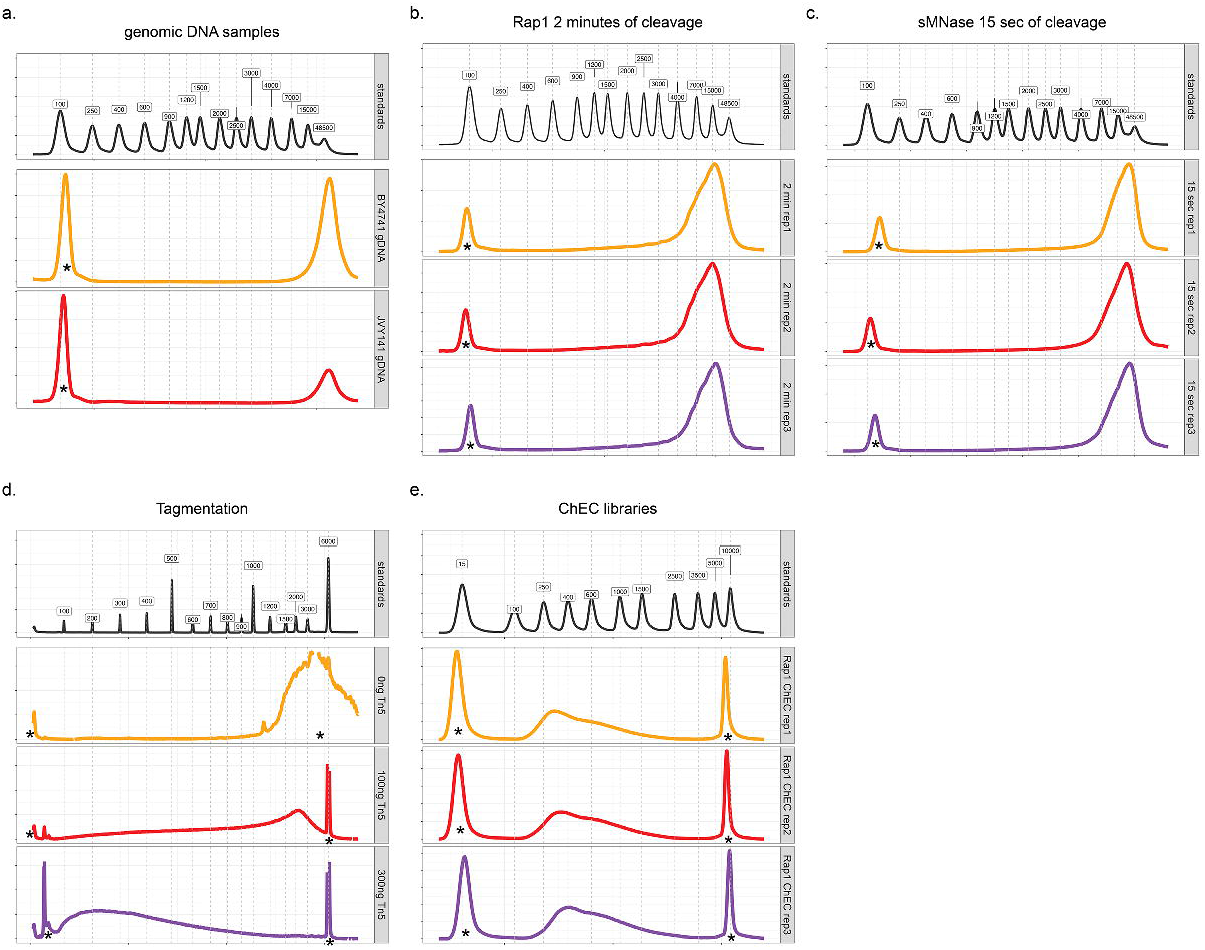
TapeStation size analysis of ChEC samples. TapeStation traces of molecular weight standards (top) or uncleaved genomic DNA isolated from two different strains (**a**), DNA from three replicates from Rap1-MNase after 2 minutes of cleavage **(b)**, DNA from three replicates of soluble MNase after 15 seconds of cleavage **(c),** DNA treated with three different Tn5 concentrations **(d)** and final amplified libraries from three different replicates of Rap1-MN ChEC **(e)**. Note: * indicates the internal size standards.

**Figure S3.**
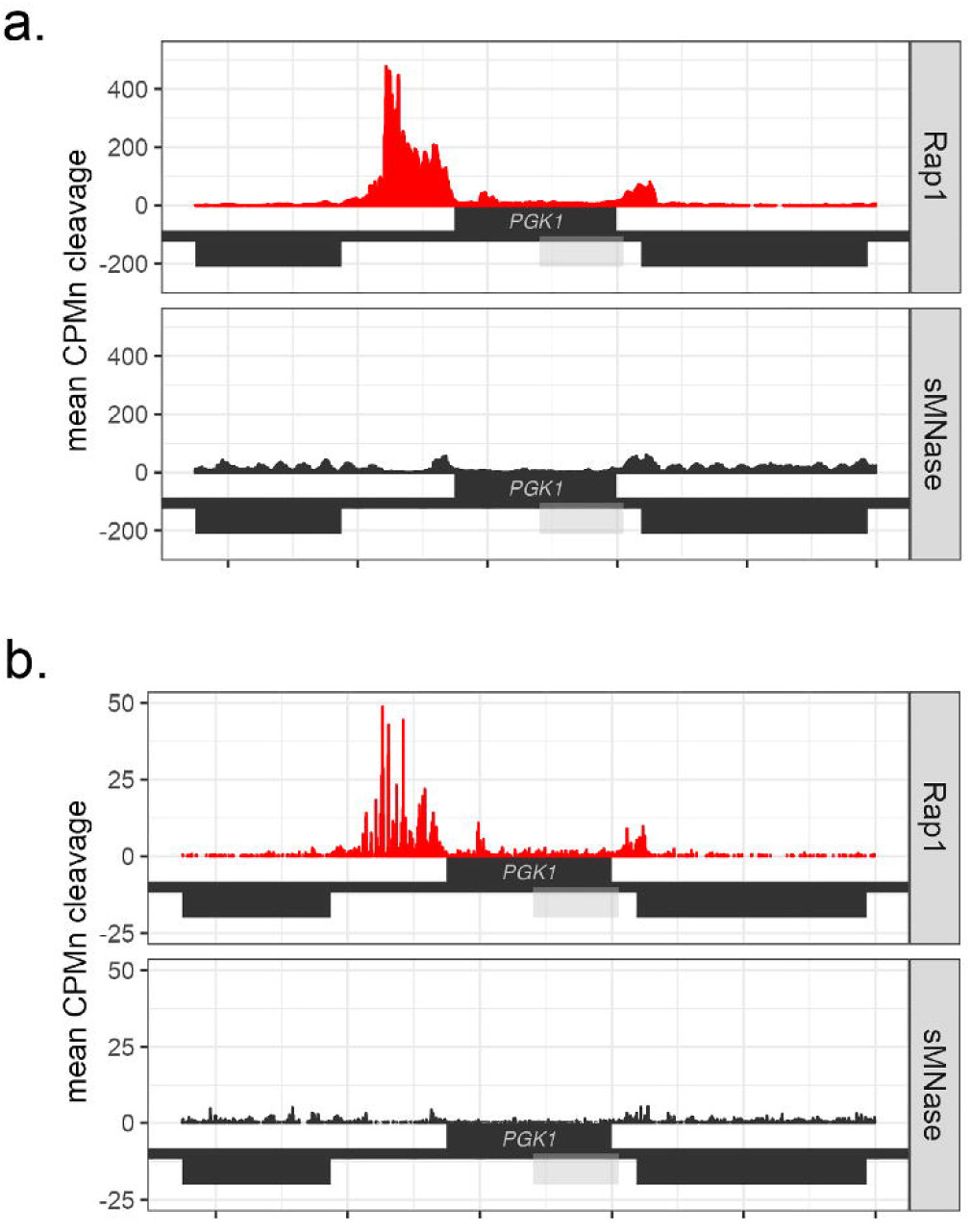
Effect of trimming reads to the first base pair on peak shape. Untrimmed (top) or trimmed (bottom; only the first base was tallied) reads from Rap1-MN ChEC-seq2 were plotted over the *PGK1* gene. Genes on the top strand are transcribed left to right and genes on the bottom strand are transcribed right to left. The light grey box indicates a dubious open reading frame (*YCR013C*).

**Figure S4.**
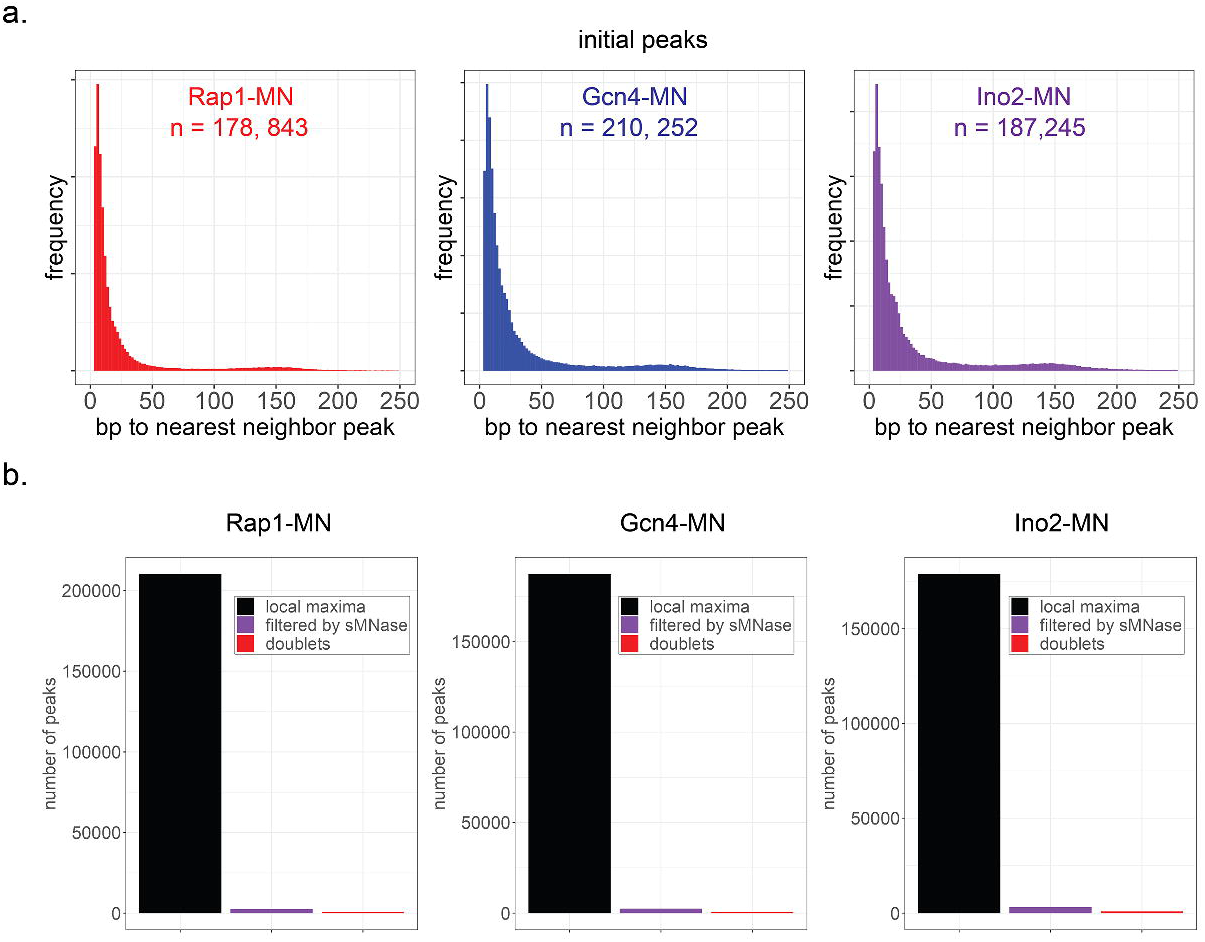
Filtering ChEC peaks to identify high-confidence sites. **(a)** The distribution of nearest distances between local maxima identified from Rap1-MN, Gcn4-MN and Ino2-MN cleavage sites. **(b)** The number of initial local maxima, sites that were significantly enriched over sMNase and doublets identified from Rap1-MN, Gcn4-MN and Ino2-MN cleavage sites.

**Figure S5.**
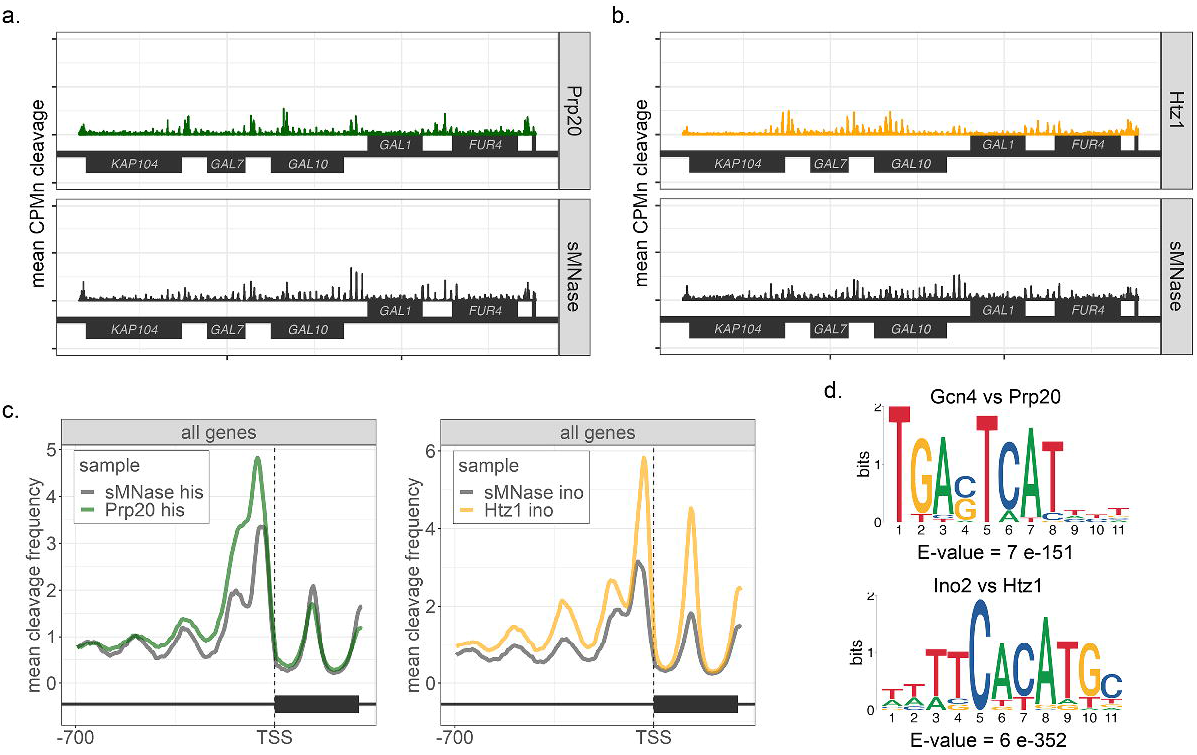
Comparison of different chromatin accessibility assays. (a) Mean cleavage frequency for Prp20 or sMNase from cells grown in SDC-histidine across the repressed *GAL1-10* locus. (b) Mean cleavage frequency for Htz1 or sMNase from cells grown in SDC-inositol across the repressed *GAL1-10* locus. (c) Meta-promoter analysis for Prp20 and sMNase from SDC-histidine (left) or Htz1 and sMNase from SDC-inositol (right). Mean CPM normalized cleavage from 700 base pairs upstream to 200 base pairs downstream of the transcriptional start site of 5348 genes with mapped transcriptional start sites (http://yeastss.org/; McMillan et al., 2019). (d) MEME analysis of high-confidence peaks identified for Gcn4 using Prp20 as a negative control (top) or Ino2 using Htz1 as a negative control (bottom).

**Figure S6.**
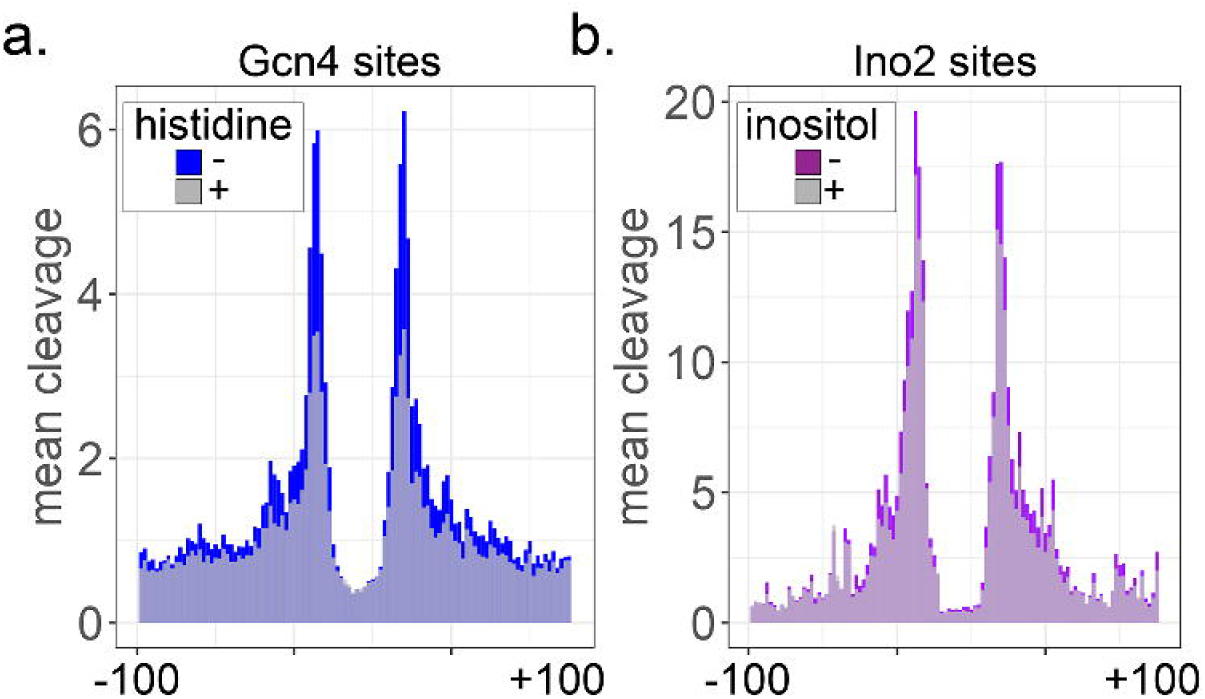
Inducible TFs show increased cleavage over their high-confidence sites. Mean cleavage by Gcn4 **(a)** or Ino2 **(b)** over high-confidence sites from cells grown in media ± histidine **(a)** or ± inositol **(b)**.

## References

Ashburner, M., C.A. Ball, J.A. Blake, D. Botstein, H. Butler, J.M. Cherry, A.P. Davis, K. Dolinski, S.S. Dwight, J.T. Eppig, M.A. Harris, D.P. Hill, L. Issel-Tarver, A. Kasarskis, S. Lewis, J.C. Matese, J.E. Richardson, M. Ringwald, G.M. Rubin, and G. Sherlock. 2000. Gene Ontology: tool for the unification of biology. Nat. Genet. 25:25–29. doi:10.1038/75556.

Bailey, T.L. 2003. Discovering Novel Sequence Motifs with MEME. Curr. Protoc. Bioinform. 00:2.4.1–2.4.35. doi:10.1002/0471250953.bi0204s00.

Balakrishnan, R., J. Park, K. Karra, B.C. Hitz, G. Binkley, E.L. Hong, J. Sullivan, G. Micklem, and J.M. Cherry. 2012. YeastMine—an integrated data warehouse for Saccharomyces cerevisiae data as a multipurpose tool-kit. Database. 2012:bar062. doi:10.1093/database/bar062.

Bergenholm, D., G. Liu, P. Holland, and J. Nielsen. 2018. Reconstruction of a Global Transcriptional Regulatory Network for Control of Lipid Metabolism in Yeast by Using Chromatin Immunoprecipitation with Lambda Exonuclease Digestion. mSystems. 3:e00215–17. doi:10.1128/msystems.00215-17.

Bolger, A.M., M. Lohse, and B. Usadel. 2014. Trimmomatic: a flexible trimmer for Illumina sequence data. Bioinformatics. 30:2114–2120. doi:10.1093/bioinformatics/btu170.

Bondra, E.R., and J. Rine. 2023. Context dependent function of the transcriptional regulator Rap1 in gene silencing and activation in Saccharomyces cerevisiae. bioRxiv. 2023.05.08.539937. doi:10.1101/2023.05.08.539937.

Brickner, J.H., and P. Walter. 2004. Gene Recruitment of the Activated INO1 Locus to the Nuclear Membrane. Plos Biol. 2:e342. doi:10.1371/journal.pbio.0020342.

Castro-Mondragon, J.A., R. Riudavets-Puig, I. Rauluseviciute, R. Berhanu Lemma, L. Turchi, R. Blanc-Mathieu, J. Lucas, P. Boddie, A. Khan, N. Manosalva Pérez, O. Fornes, T.Y. Leung, A. Aguirre, F. Hammal, D. Schmelter, D. Baranasic, B. Ballester, A. Sandelin, B. Lenhard, K. Vandepoele, W.W. Wasserman, F. Parcy, and A. Mathelier. 2021. JASPAR 2022: the 9th release of the open-access database of transcription factor binding profiles. Nucleic Acids Res. 50:gkab1113-. doi:10.1093/nar/gkab1113.

Engel, S.R., F.S. Dietrich, D.G. Fisk, G. Binkley, R. Balakrishnan, M.C. Costanzo, S.S. Dwight, B. C. Hitz, K. Karra, R.S. Nash, S. Weng, E.D. Wong, P. Lloyd, M.S. Skrzypek, S.R. Miyasato, M. Simison, and J.M. Cherry. 2013. The Reference Genome Sequence of Saccharomyces cerevisiae: Then and Now. G3: GenesGenomesGenet. 4:389–398. doi:10.1534/g3.113.008995.

Frasch, M. 1991. The maternally expressed Drosophila gene encoding the chromatin-binding protein BJ1 is a homolog of the vertebrate gene Regulator of Chromatin Condensation, RCC1. EMBO J. 10:1225–1236. doi:10.1002/j.1460-2075.1991.tb08064.x.

Ghaemmaghami, S., W.K. Huh, K. Bower, R.W. Howson, A. Belle, N. Dephoure, E.K. O’Shea, and J.S. Weissman. 2003. Global analysis of protein expression in yeast. Nature. 425:737– 41.

Gilmour, D.S., and J.T. Lis. 1984. Detecting protein-DNA interactions in vivo: distribution of RNA polymerase on specific bacterial genes. Proc Natl Acad Sci U S A. 81:4275–9.

Guillemette, B., A.R. Bataille, N. Gevry, M. Adam, M. Blanchette, F. Robert, and L. Gaudreau. 2005. Variant histone H2A.Z is globally localized to the promoters of inactive yeast genes and regulates nucleosome positioning. PLoS Biol. 3:e384.

Hennig, B.P., L. Velten, I. Racke, C.S. Tu, M. Thoms, V. Rybin, H. Besir, K. Remans, and L.M. Steinmetz. 2017. Large-Scale Low-Cost NGS Library Preparation Using a Robust Tn5 Purification and Tagmentation Protocol. G3: GenesGenomesGenet. 8:79–89. doi:10.1534/g3.117.300257.

Heyken, W.T., A. Repenning, J. Kumme, and H.J. Schuller. 2005. Constitutive expression of yeast phospholipid biosynthetic genes by variants of Ino2 activator defective for interaction with Opi1 repressor. Mol Microbiol. 56:696–707.

Hinnebusch, A.G. 1985. A hierarchy of trans-acting factors modulates translation of an activator of amino acid biosynthetic genes in Saccharomyces cerevisiae. Mol Cell Biol. 5:2349–60.

Horz, W., Werner, and Altenberg. 1981. Sequence specific cleavage of DNA by micrococcal nuclease. Nucleic Acids Research. 12:2643–2658. doi:10.1093/nar/9.12.2643.

Johnson, D.S., A. Mortazavi, R.M. Myers, and B. Wold. 2007. Genome-Wide Mapping of in Vivo Protein-DNA Interactions. Science. 316:1497–1502. doi:10.1126/science.1141319.

Langmead, B., and S.L. Salzberg. 2012. Fast gapped-read alignment with Bowtie 2. Nat. Methods. 9:357–359. doi:10.1038/nmeth.1923.

Li, H., B. Handsaker, A. Wysoker, T. Fennell, J. Ruan, N. Homer, G. Marth, G. Abecasis, R. Durbin, and 1000 Genome Project Data Processing Subgroup. 2009. The Sequence Alignment/Map format and SAMtools. Bioinformatics. 25:2078–2079. doi:10.1093/bioinformatics/btp352.

Love, M.I., W. Huber, and S. Anders. 2014. Moderated estimation of fold change and dispersion for RNA-seq data with DESeq2. Genome Biol. 15:550. doi:10.1186/s13059-014-0550-8.

McMillan, J., Z. Lu, J.S. Rodriguez, T.-H. Ahn, and Z. Lin. 2019. YeasTSS: an integrative web database of yeast transcription start sites. Database. 2019:baz048. doi:10.1093/database/baz048.

Mittal, C., M.J. Rossi, and B.F. Pugh. 2021. High similarity among ChEC-seq datasets. bioRxiv. 2021.02.04.429774. doi:10.1101/2021.02.04.429774.

Ohya, Y., N. Umemoto, I. Tanida, A. Ohta, H. Iida, and Y. Anraku. 1991. Calcium-sensitive cls mutants of Saccharomyces cerevisiae showing a Pet-phenotype are ascribable to defects of vacuolar membrane H(+)-ATPase activity. J. Biol. Chem. 266:13971–7.

Raisner, R.M., P.D. Hartley, M.D. Meneghini, M.Z. Bao, C.L. Liu, S.L. Schreiber, R, O.J. o, and H.D. Madhani. 2005. Histone variant H2A.Z marks the 5’ ends of both active and inactive genes in euchromatin. Cell. 123:233–48.

Rhee, H.S., and B.F. Pugh. 2011. Comprehensive genome-wide protein-DNA interactions detected at single-nucleotide resolution. Cell. 147:1408–19. doi:10.1016/j.cell.2011.11.013.

Rossi, M.J., W.K.M. Lai, and B.F. Pugh. 2017. Correspondence: DNA shape is insufficient to explain binding. Nat. Commun. 8:15643. doi:10.1038/ncomms15643.

Schmid, M., T. Durussel, and U.K. Laemmli. 2004. ChIC and ChEC; genomic mapping of chromatin proteins. Mol Cell. 16:147–57.

Shore, D., S. Zencir, and B. Albert. 2021. Transcriptional control of ribosome biogenesis in yeast: links to growth and stress signals. Biochem. Soc. Trans. 49:1589–1599. doi:10.1042/bst20201136.

Skene, P.J., and S. Henikoff. 2017. An efficient targeted nuclease strategy for high-resolution mapping of DNA binding sites. eLife. 6:e21856. doi:10.7554/elife.21856.

Solomon, M.J., P.L. Larsen, and A. Varshavsky. 1988. Mapping protein-DNA interactions in vivo with formaldehyde: evidence that histone H4 is retained on a highly transcribed gene. Cell. 53:937–47.

Wagner, C., M. Dietz, J. Wittmann, A. Albrecht, and H.J. Schuller. 2001. The negative regulator Opi1 of phospholipid biosynthesis in yeast contacts the pleiotropic repressor Sin3 and the transcriptional activator Ino2. Mol Microbiol. 41:155–66.

Wu, T., E. Hu, S. Xu, M. Chen, P. Guo, Z. Dai, T. Feng, L. Zhou, W. Tang, L. Zhan, X. Fu, S. Liu, X. Bo, and G. Yu. 2021. clusterProfiler 4.0: A universal enrichment tool for interpreting omics data. Innov. 2:100141. doi:10.1016/j.xinn.2021.100141.

Yu, G., L.-G. Wang, Y. Han, and Q.-Y. He. 2012. clusterProfiler: an R Package for Comparing Biological Themes Among Gene Clusters. *OMICS: A J*. Integr. Biol. 16:284–287. doi:10.1089/omi.2011.0118.

Zentner, G.E., S. Kasinathan, B. Xin, R. Rohs, and S. Henikoff. 2015. ChEC-seq kinetics discriminates transcription factor binding sites by DNA sequence and shape in vivo. Nat Commun. 6:8733. doi:10.1038/ncomms9733.

