## Supplementary material for "ChEC-seq2: an improved Chromatin Endogenous Cleavage sequencing method and bioinformatic analysis pipeline for mapping *in vivo* protein-DNA interactions": Detailed ChEC-seq2 protocol

**Chromatin endogenous cleavage protocol**

### Chromatin digestion

1. Grow cells in 10ml overnight at 30°C, 200 rpm.
2. Dilute cultures into 50ml media to $OD_{600}$ ~ 0.1.
3. When cultures reach $OD_{600}$ = 0.5 - 0.8, harvest 25 ODs (*i.e.* 50ml if the OD600 = 0.5) of cells by centrifugation at 2500 x *g* for 1 minute.
4. Resuspend cells in 1 ml Buffer A and transfer to a 1.5 ml tube.
5. Pellet cells by centrifugation at 2500 x *g* for 1 minute, remove supernatant.
6. Wash cells 2 x 1 ml Buffer A, removing supernatant.
7. Resuspend cells in 600 µl Buffer A + 0.1% Digitonin.
8. Transfer tube to a 30°C heat block and incubate for 5 minutes.
9. Add 5 µl of 333 mM $CaCl_2$, mix by inverting, incubate at 30°C for the appropriate cleavge time (determined empirically for each protein).
10. To stop the reaction, remove 200 µl cells and combine with 200 µl 2x Stop Buffer.
11. Add 8 µl Proteinase K (20 µg/µl) and mix.
12. Incubate at 50°C, agitating 800 rpm for 30 minutes in a thermomixer.
13. Remove samples from thermomixer and cool at room temperature for 5 minutes.
14. Add 400 µl Phenol-Chloroform-Isoamyl Alcohol (25:24:1), pH 7.8, mix.
15. Centrifuge at 24,000 x *g*, 5 minutes.
16. Transfer aqueous phase to a phase-lock tube.
17. Add 200 µl Phenol-Chloroform-Isoamyl Alcohol (25:24:1), pH 7.8.
18. Invert 10x to mix.
19. Centrifuge at 24,000 x *g* for 5 minutes.
20. Transfer aqueous phase to a tube containing 1 ml 100% Ethanol.
21. Add 2 µl of linear acrylamide (5 µg/µl).
22. Invert 10x to mix.
23. Incubate at -80°C for 30 minutes.
24. Centrifuge at 24,000 x g, 4°C for 10 minutes.
25. Pour off supernatant.
26. Wash DNA pellet in 1 ml of 70% ethanol.
27. Centrifuge at 24,000 x g for 1 minute.
28. Pour off supernatant. Collect residual ethanol by centrifugation and remove by pipetting.
29. Dry DNA pellet until all ethanol had evaporated.
30. Add 58 µl of 10 mM Tris-HCl, pH 8.5 to DNA pellet.
31. Incubate overnight at room temperature.
32. Incubate at 37°C for 30 minutes.
33. Add 2 µl RNase A (10 µg/µl) to DNA.
34. Incubate at 37°C for 30 minutes.
35. Evaluate molecular weight of DNA by gel electrophoresis, 0.8% agarose or TapeStation.
36. Quantify DNA concentration with the Qubit double-stranded DNA, Broad Range Assay.
37. Stored DNA at 4°C until library preparation was performed (up to a month), then stored at -20°C.

### <u>Buffer A</u>

| | |
|---|---|
| 15 mM | Tris-HCl, pH 7.5 |
| 80 mM | KCl |
| 0.1 mM | EGTA |
| 1.0 mM | PMSF |
| 0.5 mM | Spermidine |
| 0.2 mM | Spermine |

-Add 1 EDTA-Free Protease Inhibitor Tab per 50 ml Buffer A (Roche; Sigma # 11873580001)

**2x Stop Buffer**

| | |
|---|---|
| 400 mM | NaCl |
| 20 mM | EDTA |
| 4 mM | EGTA |

**Library Preparation & Sequencing**

**End repair and ligation**

1. Dilute each DNA sample to 10 ng/µl x 30 µl in PCR tubes.
2. For each sample, in a 0.2ml PCR tube, combine:
   2.5 µl 10x CutSmart Buffer
   1.5 µl water
   20µl gDNA (10ng/µl)
   1µl Quick CIP (NEB #M0508)
3. Mix carefully and then spun tubes to collect droplets.
4. Heat in a thermocycler:
   - 37°C - 10:00
   - 80°C - 5:00
   - 4°C - HOLD
5. Extend DNA ends and add a 5' phosphate, add 10µl of the following to each reaction:
   - 1 µl 10x CutSmart Buffer (NEB #E1201)
   - 3.5 µl 1 mM dNTPs (NEB #E1201)
   - 4.5 µl water
   - 1µl Blunting Enzyme Mix (NEB #E1201)
6. Heat in a thermocycler:
   - 25°C - 30:00
   - 70°C - 10:00
   - 4°C - HOLD
7. Ligate i5 Y-Adapter to cleaved, repaired gDNA (NEB #M2200). To each reaction, add:
   - 32.5 µl 2x Quick Ligation Buffer (NEB #M2200)
   - 0.5 µl 45 µM i5 Y-Adapter (NEB #M2200)
   - 2 µl Quick Ligase (NEB #M2200)
   - 32.0 µl water
   - Total Volume: 100 µl
8. Heated in a thermocycler:
   - 25°C - 20:00
   - 4°C - HOLD
9. Allow ligation reactions to come to room temperature.
10. Add 70 µl SPRIselect beads (Final [Beads] = 0.7x) to each 100 µl ligation reaction (PCR strip tubes) and mix by pipetting 20x.
11. Incubate at room temperature for 1 minute,
12. Place PCR tubes onto magnetic stand and waited for beads to adhere, carefully remove supernatant.
13. Add 180 µl of 85% ethanol, waited 30 seconds, and then remove ethanol.
14. Repeat ethanol wash and carefully remove any residual ethanol.
15. Air-dry beads for 3 minutes.
16. Remove tube from rack and resuspend beads in 25 µl of water.
17. Incubate at room temperature for 1 minute, then place tube back onto magnetic stand and wait for beads to adhere.
18. Collect DNA-containing supernatant and transfer to new PCR tubes.
19. Quantify DNA concentration with 2 µl using Qubit 1x High Sensitivity dsDNA kit.
20. Store purified, ligated DNA at 4°C overnight.

**Tagmentation and library amplification**

21. To 10 ng of ligated ChEC-gDNA in a 24 µl reaction volume, add:
    - 6 µl 4x Tagmentation Buffer (see below)
    - 6 µl 100% DMF (dimethylformamide)
    - ~ µl 10 ng DNA
    - 1 µl Tn5 (300ng), loaded with i7-A
    - Water to 24 µl
22. Mix reactions by flicking wells, and then spin plate to collect droplets.
23. In a thermocycler, heat at 55°C for 10 minutes, then hold at 10°C
24. Add 16.8 µl SPRIselect beads to each 24 µl Tagmentation reaction (Final [Beads] = 0.7x) in a PCR tube.
25. Mix by pipetting 20x.
26. Incubate at room temperature for 1 minute
27. Place tubes onto magnetic stand and wait for beads to adhere
28. Carefully remove supernatant
29. Add 180 µl of 85% ethanol, wait 30 seconds, and then removed ethanol.
30. Repeat ethanol wash and remove any residual ethanol.
31. Air-dry beads for 3 minutes
32. Remove tubes from rack and resuspend beads in 10 µl of water.
33. Incubate at room temperature for 1 minute, then place tube back onto magnetic stand and wait for beads to adhere.
34. Collect supernatant and transfer to new PCR tubes.
35. Prepare library amplification reaction:
    - 6 µl 2x KAPA Library Amplification Master Mix
    - 0.5 µl 10 µM S502 Index-Primer ([Final] = 0.42 µM)
    - 0.5 µl 10 µM N7xx Index-Primer ([Final] = 0.42 µM)
    - 5 µl Purified, Tagmented DNA
    - Total Volume = 12 µl
36. Mix reactions and spin to collect droplets.
37. Transfer tubes to a thermocycler and run the following protocol:
    - 72°C - 3:00
    - 98°C - 2:45
    - 98°C - 0:15*
    - 62°C - 0:30* *15 cycles
    - 72°C - 1:30*
    - 72°C - 3:00
    - 4°C - HOLD
38. Add 8.4 µl of SPRIselect beads directly to PCR tubes containing 12 µl of Library Amplification Reaction (Final [Beads] = 0.7x).
39. Mix by pipetting 20x.
40. Incubate at room temperature for 1 minutes
41. Place PCR strip tubes onto magnetic stand and wait for beads to adhere
42. Carefully remove supernatant
43. Add 180 µl of 85% ethanol, wait 30 seconds, and then remove ethanol.
44. Repeat ethanol wash and remove any residual ethanol.
45. Air-dry beads for 3 minutes
46. Remove tubes from rack and resuspend beads in 10 µl of water.
47. Incubate at room temperature for 1 minute, then place tube back onto magnetic stand and wait for beads to adhere.
48. Collect DNA-containing supernatant and transfer to new PCR tubes.

*49.* Quantify DNA concentration using the QuBit dsDNA High Sensitivity Assay (see table)
*50.* Combine 2 µl sample with 190 µl Buffer to evaluate molecular weight of libraries on the TapeStation with the D5000 kit.
*51.* Pool sample libraries with different indexes
*52.* Combine equimolar amounts of each library into a 1.5 ml tube.
*53.* The pooled library concentration should be 2 nM in 100µl.
